# Intramammary infection of bovine H5N1 influenza virus in ferrets leads to transmission in suckling neonates

**DOI:** 10.1101/2024.11.15.623885

**Authors:** PH Baker, M Moyer, Y Bai, LS Stafford, C Lee, AA Kelvin, SN Langel

## Abstract

The spread of H5N1 clade 2.3.4.4b in dairy herds raises concerns about zoonotic transmission due to its high viral load in milk, a key contact point between livestock and humans. H5N1 clade 2.3.4.4b exhibits tropism for the mammary gland, with milk from infected animals containing high levels of infectious virus, posing potential risks to offspring via breastfeeding. Using a lactating ferret model, we demonstrate that mammary gland infection with bovine H5N1 transmits the virus to suckling kits, resulting in neonatal mortality. Viral RNA levels increased in milk and remained high in mammary tissue, with infected kits exhibiting elevated viral RNA in the oral and nasal cavities and feces. Additionally, we detected the H5N1 receptor, α2,3 sialic acid, in ferret and human mammary tissue. These data demonstrate that H5N1 clade 2.3.4.4b infection in lactating dams leads to mastitis-related disease and transmits to suckling pups, resulting in mortality among neonates.

## Introduction

Highly pathogenic avian influenza (HPAI) H5N1 clade 2.3.4.4b was first detected in the U.S. in late 2021, causing widespread outbreaks in wild bird populations and commercial poultry, resulting in high mortality^1^. H5N1 clade 2.3.4.4b has spilled over into an unprecedented number of mammalian species worldwide^2^, heightening concerns about its potential pandemic risk to humans. In March 2024, H5N1 clade 2.3.4.4b was detected in dairy cows for the first time^3,4^, showing a marked tropism for the mammary gland^5^. As of November 21, dairy cows from 15 states have been affected, with 549 farms reporting cases to the CDC^6^. In connection with this outbreak, 53 confirmed human cases have been reported: 31 linked to cattle, 21 associated with infected poultry, and one with an unidentified source^6^. Human cases are primarily among farm workers with mild to moderate respiratory symptoms and conjunctivitis, highlighting a serious public health concern.

Transmission of the virus among cattle and within dairy herds is believed to occur primarily through the interstate movement of infected animals and direct contact via contaminated milk and milking equipment^7^. Notably, H5N1 clade 2.3.4.4b has shown a pronounced affinity for the bovine mammary gland. Mammary tissue and milk of infected animals contain markedly elevated levels of viral RNA and infectious virus^5,8,9^. Infected cows were initially identified following reports of reduced milk yields on farms and exhibited nonspecific disease signs, including decreased rumination, abnormal milk characteristics, and respiratory distress^5^. Influenza A virus has been rarely documented in cattle, but previous studies have shown evidence of its associated with reductions in milk yield in dairy cows^10,11^ and direct replication in the ferret mammary gland^12^.

The potential for a respiratory virus like H5N1 to infect a non-traditional site, such as the mammary gland, raises concerns about milk as a possible interface for virus transmission between infected livestock and humans. Understanding the mechanisms that govern H5N1 infection, transmission, and immune protection within the mammary gland will be crucial to controlling the current outbreak and preventing future ones in both humans and animals. This will also aid in understanding the risk of infection and disease in suckling neonates, a population that remains understudied in the context of H5N1 infection. Investigating H5N1 infection in the mammary gland through controlled studies in cattle is challenging and labor-intensive, primarily due to the complexities of working with large animals in biosafety level 3 (BSL-3) containment. Therefore, developing preclinical models of H5N1 2.3.4.4b infection and transmission in the mammary gland via milk is essential to enhance our understanding of these unique infections. Ferrets are the gold-standard preclinical model for studying influenza virus susceptibility, pathogenesis, and transmission. Our previous research demonstrated that the lactating ferret mammary gland can be infected with the H1N1 influenza virus^12^. This study aimed to evaluate the potential for H5N1 clade 2.3.4.4b transmission through the milk of lactating dams and to assess its pathogenesis in suckling offspring.

## Results

### Intramammary bovine H5N1 inoculation causes severe morbidity and mortality in ferret dams and kits

To investigate bovine H5N1 virus infection in the mother-infant dyad, we inoculated mammary glands of lactating ferrets at 2.5 weeks postpartum with A/bovine/Ohio/B24OSU-342/2024 H5N1 clade 2.3.4.4b. Lactating mother ferrets (dams) were housed and left to nurse their 2.5-week-old kits (n=3 kits/dam) after inoculation. Lactating mammary glands were inoculated at 10^5^ EID_50_ (3 glands/ferret) and animals were monitored and sampled up to 6 days post-inoculation (DPI) (**Figure 1A**). Mammary glands were also monitored for signs of mastitis post-inoculation. Intramammary H5N1 clade 2.3.4.4b virus infection resulted in 100% mortality of kits by 4 DPI and in dams by 6 DPI (**Figure 1B**). Ferret dams showed signs of lethargy, reduced feed intake, and weight loss. Weights significantly decreased by 6 DPI (*P* < 0.0001) and dams were euthanized on 6 DPI due to a humane endpoint weight threshold (**Figure 1C**). Ferret kits exposed to H5N1-positive milk experienced weight gain (*P* = 0.02) from 0 to 2 DPI; however, experienced significant weight loss from 2 to 3 DPI (*P* = 0.0005) and 3 to 4 DPI (*P* = 0.04; **Figure 1D**). Intramammary H5N1 infection in the dams caused illness, with a high fever onset at 2 DPI that persisted throughout the experimental period (**Figure 1E**). All kits exhibited a slight temperature increase, with some developing a fever by 2 DPI; however, temperatures in all kits returned to baseline by 3 DPI (**Figure 1F**). The remaining kit showed a decrease in temperature and was hypothermic by the time of humane euthanasia at 4 DPI. Notably, at 4 DPI, we observed a decrease in milk production. In one ferret dam, an infected mammary gland displayed abnormal, thick, colostrum-like milk, while the other glands from both ferrets produced watery secretions, indicating signs of mastitis.

**Figure 1:**
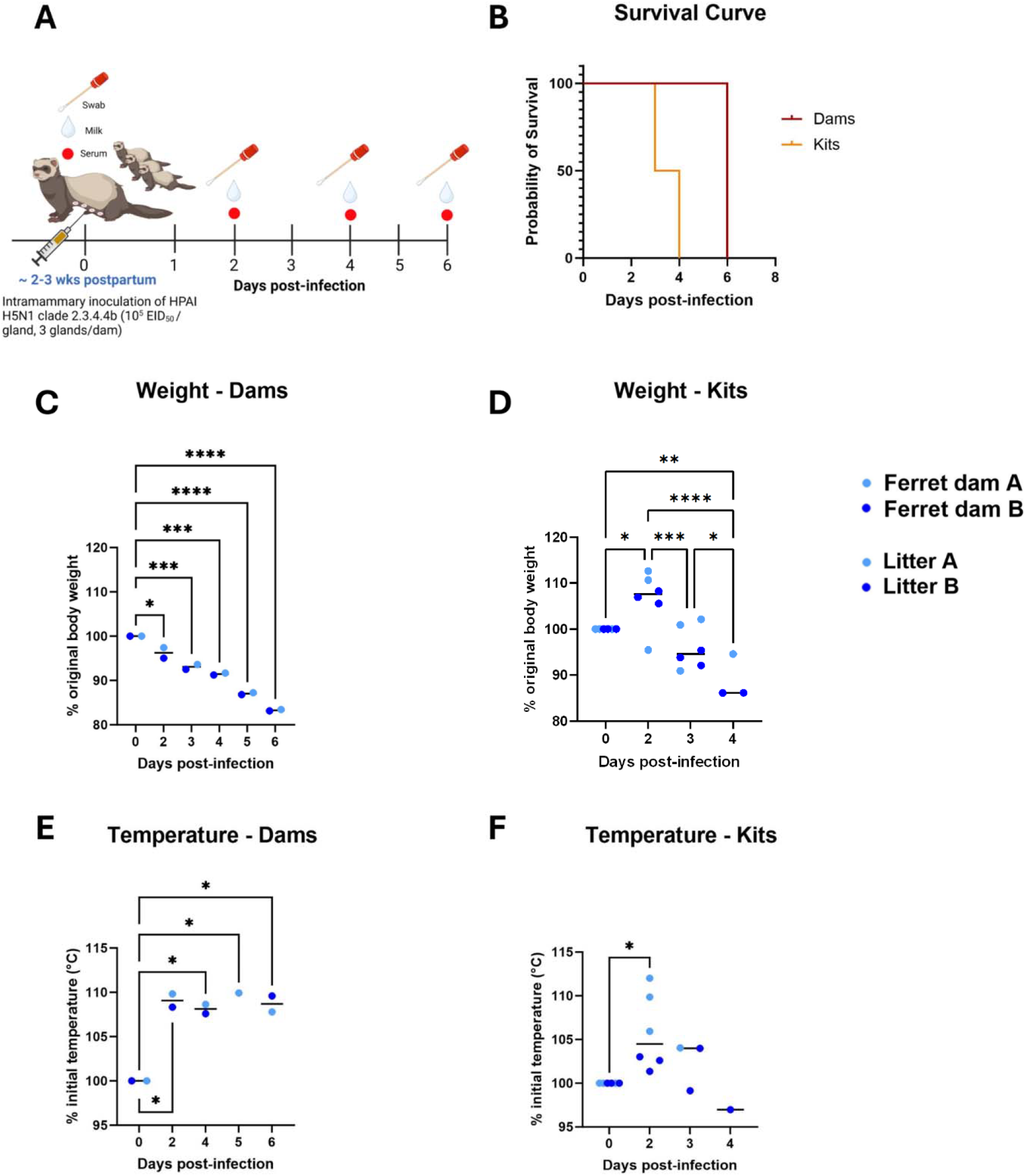
Intramammary inoculation of bovine H5N1 results in 100% mortality in ferret dams and kits. **(A)** Experimental design for intramammary infection in lactating ferret dams. Lactating dams were inoculated in the mammary gland with HPAI H5N1 clade 2.3.4.4b (10^5^ EID_50_/gland) and monitored for survival up to 6 days post-infection (DPI). (**B)** Survival curve for dams and kits. Weight **(C, D)** and temperature **(E, F)** were measured for dams and kits up to 6 DPI. Each point represents an individual dam or kit measurement. Asterisks indicate statistically significant differences (\**P* ≤ 0.05, \*\**P* ≤ 0.01, \*\*\**P* ≤ 0.001, \*\*\*\**P* ≤ 0.0001).

### Intramammary inoculation of bovine H5N1 virus induces viral persistence in milk and the mammary gland

Milk was collected from lactating dams prior to infection and at 2, 4 and 6 DPI. H5N1 viral RNA levels in milk showed an increase at 2 DPI (*P* = 0.07) and 6 DPI (*P* = 0.06) compared to baseline (**Figure 2A**). The presence of virus in the mammary gland suggests active viral replication within the gland, accounting for the observed increase in viral levels compared to 0 DPI. At necropsy, one uninfected mammary gland was collected from each ferret dam as a control, confirming it remained uninfected in contrast to the inoculated glands (**Figure 2B**).

**Figure 2:**
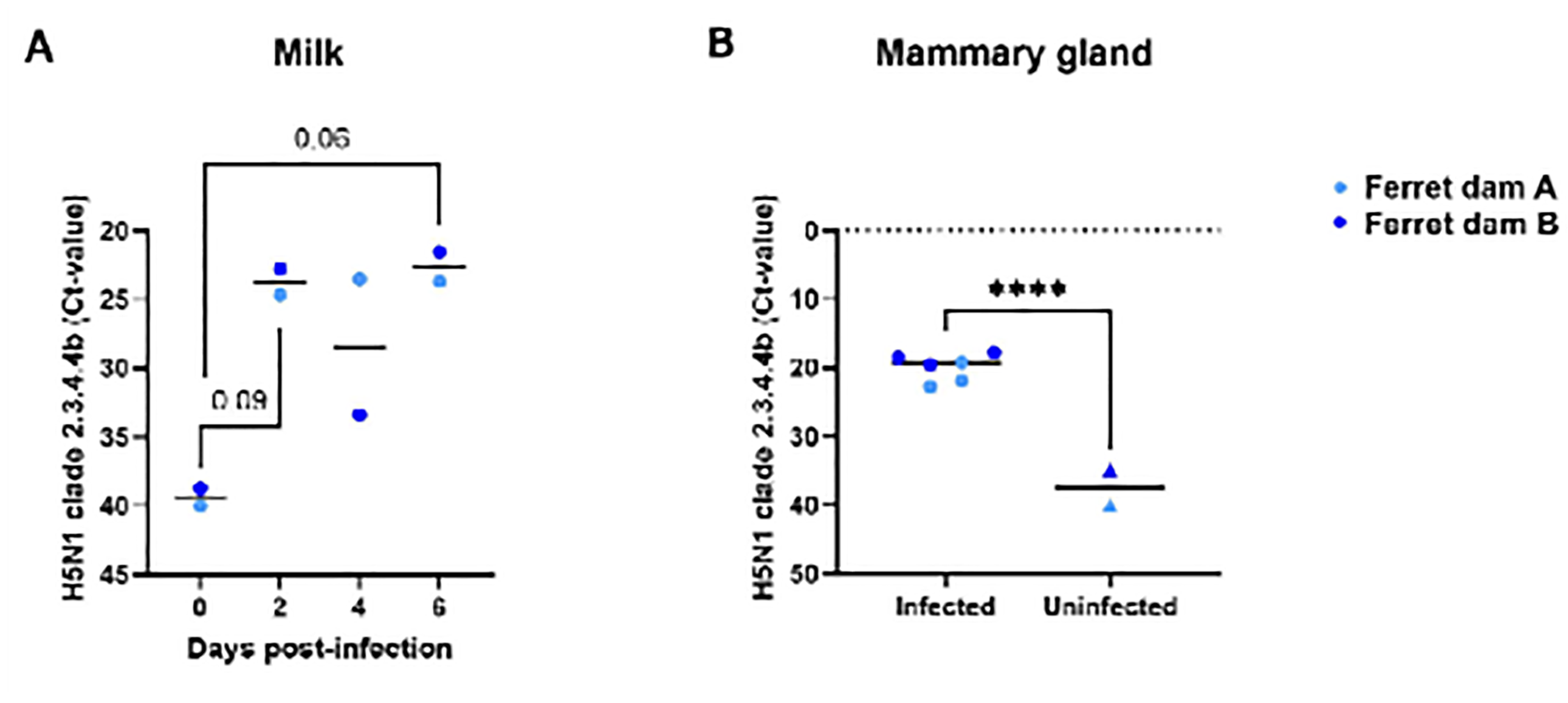
High levels of viral RNA present in milk and mammary glands of lactating ferret dams after intramammary bovine H5N1 inoculation. Cycle threshold (Ct) values were determined by RT-qPCR detection in milk from each dam **(A)** and inoculated (n=3 glands/dam) and uninoculated dam mammary glands (n=1 gland/dam). **(B)** All values resulting in an undetermined RT-qPCR value were assigned a Ct value of 40. Asterisks represent statistically significant differences between parameters (\**P* ≤ 0.05, \*\**P* ≤ 0.01, *** *P* ≤ 0.001, \*\*\*\**P* ≤ 0.0001).

### Bovine H5N1 virus is transmitted by milk, leading to viral transmission in the oral cavity and upper respiratory tract of suckling kits

Oral and nasal swabs were collected from all animals at baseline and post-intramammary inoculation. There were no differences in viral RNA from oral swabs collected from ferret dams throughout the sampling period (*P* ≥ 0.6; **Figure 3A**). Interestingly, there was a significant increase in viral RNA in oral swabs from suckling kits from 0 to 2 (*P* < 0.0001) and 4 DPI (*P* < 0.0001) and from 2 to 4 DPI (*P* = 0.0004), with viral RNA reaching peak concentrations at 4 DPI (**Figure 3B**), consistent with viral transmission in milk.

**Figure 3:**
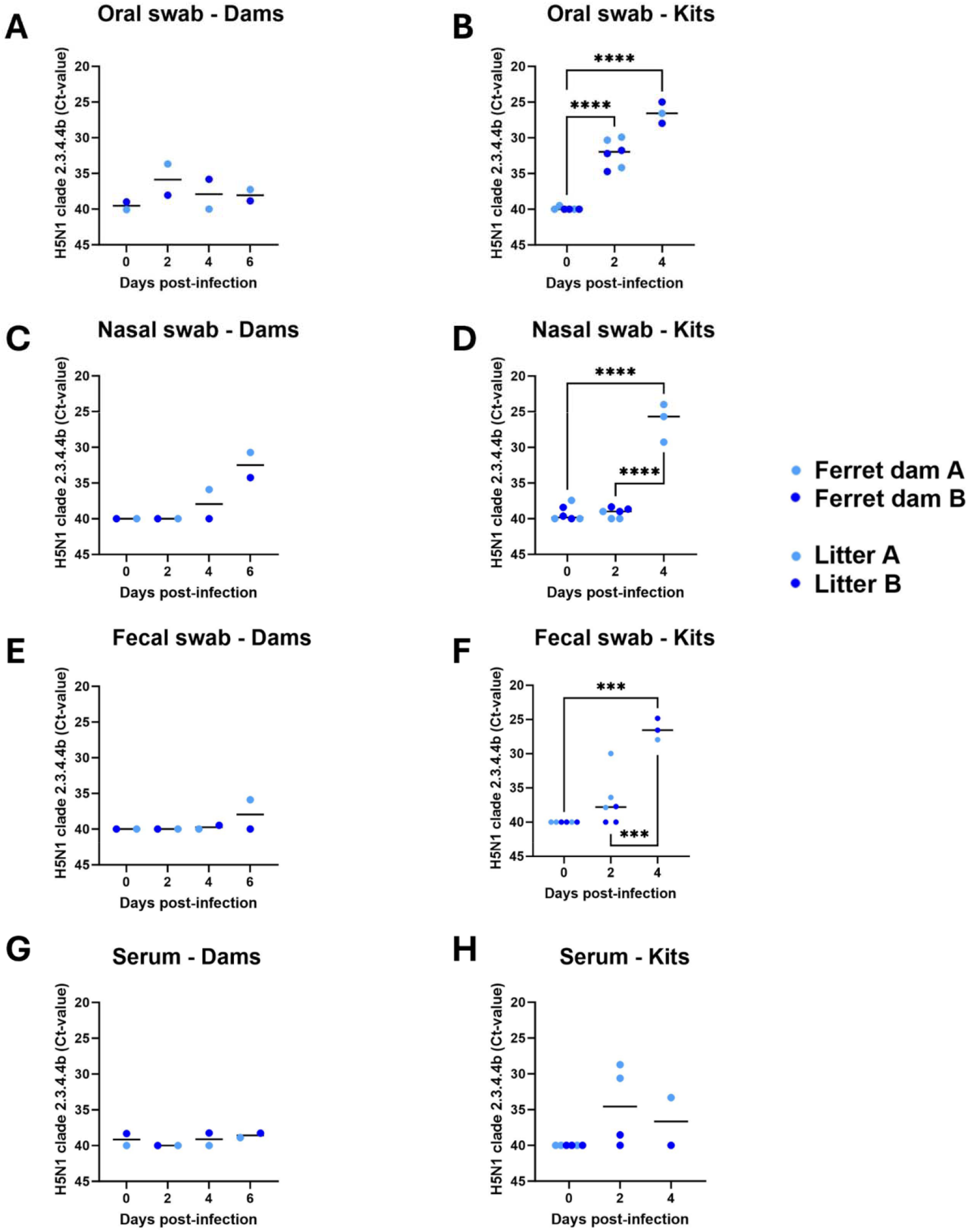
Viral RNA detection in oral, nasal, fecal swabs and serum collected from ferret dams and kits following intramammary inoculation of bovine H5N1 virus. Ct values were determined by RT-qPCR in dam and kit oral swabs (**A, B**), dam and kit nasal swabs (**C, D**), dam and kit fecal swabs (**E, F)** and dam and kit serum (**G, H**) up to 6 days post infection. All values resulting in an undetermined RT-qPCR value were assigned a Ct value of 40. Asterisks represent statistically significant differences between parameters (\**P* ≤ 0.05, \*\**P* ≤ 0.01, \*\*\**P* ≤ 0.001, \*\*\*\**P* ≤ 0.0001).

Nasal swabs collected from ferret dams at 0 and 2 DPI showed no indication of H5N1 viral RNA (*P* > 0.99). However, after prolonged contact with suckling kits, there was an increase in viral RNA in nasal swabs from 0 to 6 DPI (*P* = 0.05) and 2 to 6 DPI (*P* = 0.05) in ferret dams (**Figure 3C**). Nasal swabs collected from suckling kits showed a significant increase in viral RNA from baseline to 4 DPI (*P* < 0.0001) and between 2 and 4 DPI (*P* < 0.0001) (**Figure 3D**).

There were no differences in fecal viral RNA measured in ferret dams following intramammary infection (*P* ≥ 0.99; **Figure 3E**). Unlike dams, kits experienced a significant increase in fecal viral RNA from 0 to 4 DPI (*P* = 0.0002) and from 2 to 4 DPI (*P* = 0.0009), suggesting viral shedding within the gastrointestinal tract through the ingestion of H5N1 positive milk. This result corresponds with the oral swab data of the kits, as viral RNA reached peak concentrations by 4 DPI, coinciding with the high levels of viral RNA being shed in feces at this time point. (**Figure 3F**). Intramammary inoculation does not result in systemic infection in ferret dams as serum samples from dams showed no differences in viral RNA at any timepoints post-intramammary inoculation (*P* ≥ 0.26; **Figure 3G**). However, serum samples collected from litter A kits (n=3) were between 28 and 34 Ct while litter B kits (n=3) remained above 38 Ct. (**Figure 3H**)

### Elevated viral burden in mammary glands of ferret dams compared to other tissues

To determine tissue tropism after intramammary inoculation, several tissues were collected at necropsy and homogenized for viral RNA via quantitative real-time PCR (RT-qPCR). As expected, the mammary glands in dams exhibited the highest levels of viral RNA compared to other tissue sites (**Figure 4A**), while only low levels were present in the upper and lower respiratory tracts. In kits surviving to 4 DPI, viral RNA was also detected in their stomach contents. In a kit sampled at 4 DPI, high viral RNA titers were detected in the bronchoalveolar lavage (BAL) fluid and lungs, whereas lower titers were observed in the soft palate and nasal turbinates (**Figure 4B**).

**Figure 4:**
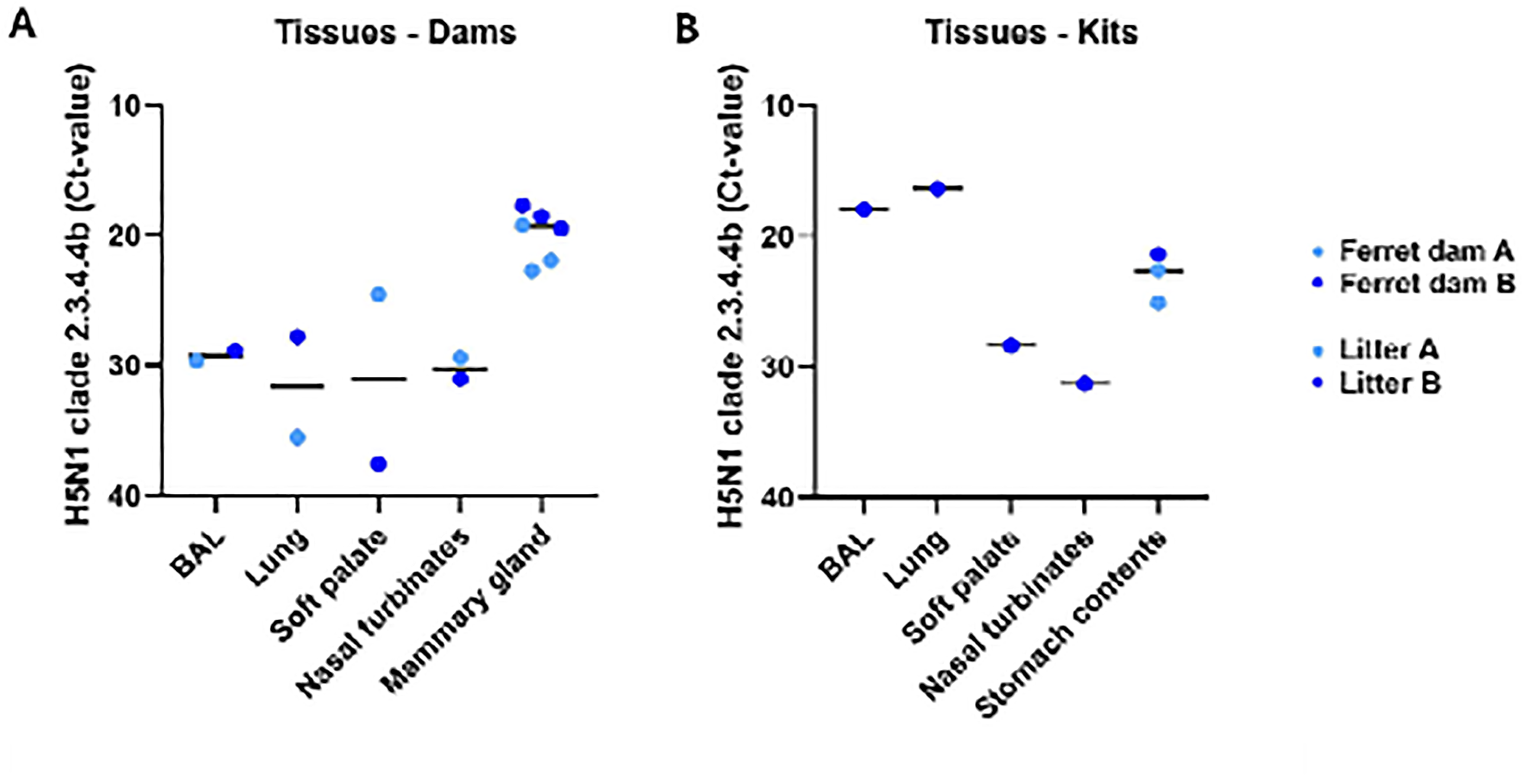
Tissue distribution of viral RNA following intramammary inoculation of bovi H5N1. Cycle threshold (Ct) values were determined by RT-qPCR in dam (**A**) and kit (**B**) bronchoalveolar lavage fluid (BAL), lung, soft palate, nasal turbinates, and mammary gland at the necropsy timepoint. All values resulting in an undetermined RT-qPCR value were assigned a Ct value of 40. Asterisks represent statistically significant differences between parameters (\**P* ≤ 0.05, \*\**P* ≤ 0.01, *** *P* ≤ 0.001, \*\*\*\**P* ≤ 0.0001).

### Ferret and human mammary gland tissue expresses the receptor for HPAI H5N1

Lactating ferret and non-lactating healthy human mammary gland tissues were stained for the expression of sialic acid with an α2,6-galactose linkage (SAα2,6-gal) and SAα2,3-gal-β(1–3)GalNAc using plant-derived lectins SNA (*Sambucus nigra*) and MAA (*Maackia amurensis* lectin II), respectively (**Figure 5**). Both ferret and human mammary tissue expressed SAα2,6-gal and SAα2,3-gal. In the lactating ferret mammary tissue, SAα2,6-gal predominately localized within the alveolar lumens and polarized to the apical membrane of mammary epithelial cells (**Figure 5A**) while SAα2,3-gal primarily located in the stromal compartment between lobules (**Figure 5B**). In the non-lactating human mammary tissue, SAα2,6-gal is primarily localized to the mammary epithelium (**Figure 5C**) while SAα2,3-gal was observed in both the epithelium and stromal compartment (**Figure 5D**).

**Figure 5:**
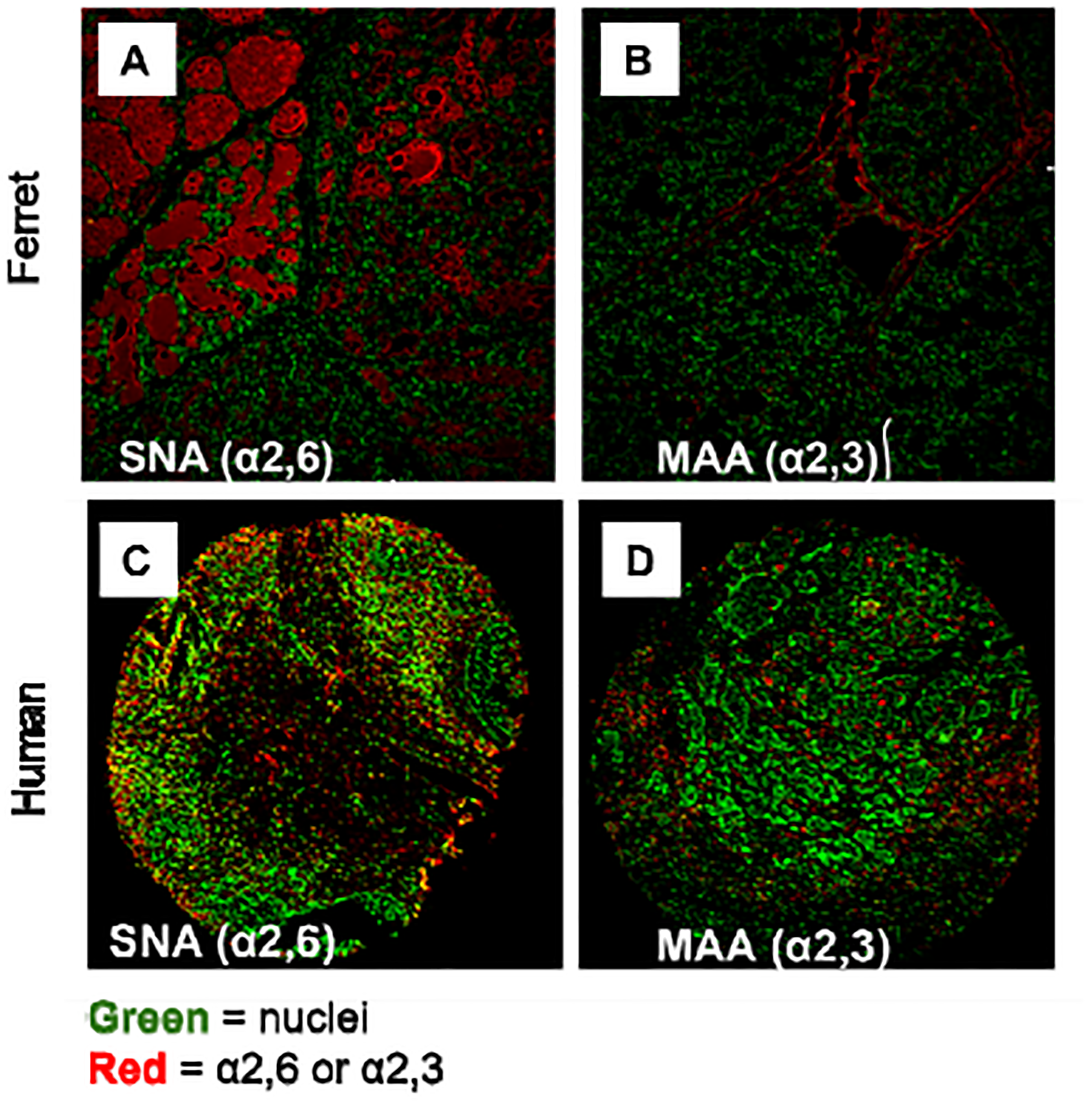
Ferret lactating and human non-lactating healthy mammary glands express both SAα2,6-gal and SAα2,3-gal. Lactating ferret (**A-B**) and non-lactating human (**C-D**) mammary tissues were incubated with fluorescently labeled plant-derived lectins SNA (*Sambucus nigra*) (**A, C**) and MAA (*Maackia amurensis* lectin II) (**B, D**) to determine expression of SAα2,6-gal and SAα2,3-gal, respectively. All cells were stained with SYTOX Green nucleic acid stain.

## Discussion

A novel HPAI H5N1 virus strain, isolated from bovine hosts, demonstrates a concerning ability to replicate in the lactating mammary gland, highlighting a risk for transmission to neonates via breastfeeding^8,9^. Our study shows that intramammary inoculation with H5N1 clade 2.3.4.4b in lactating ferrets results in transmission to suckling neonates, leading to 100% mortality in both dams and kits. High levels of viral RNA were detected in the mammary glands of ferret dams, and viral RNA was shed in milk and transmitted to the suckling kit respiratory tract. Notably, we observed that SA α2,3-gal is expressed in both lactating ferret and non-lactating human mammary glands. These findings highlight a novel and concerning route of potential zoonotic transmission through milk or contact with infected animals, posing a potential risk for lactating individuals and their infants.

Previous work by Paquette et al. demonstrated that intramammary inoculation with H1N1 led to infection in both ferret dams and kits, causing 100% mortality in suckling kits^12^. In this study, H1N1-infected kits experienced a 70% mortality rate by 7 DPI, while H5N1-infected kits reached 100% mortality by 4 DPI. In contrast, intramammary H1N1-infected dams showed a temperature spike at 2 DPI and a 10% weight reduction over seven days. H5N1-infected dams exhibited more severe symptoms, including weight reduction and a higher temperature increase. In both studies, viral kinetics in the oral and nasal cavities suggest that kits developed respiratory infections directly from H5N1-positive milk, leading to respiratory transmission to the mother. These findings indicate that intramammary H5N1 infection causes more severe disease in lactating ferrets and their kits than H1N1.

To date, there are two reports documenting experimental intramammary H5N1 infection in lactating dairy cows^8,9^. Halwe et al. reported mortality outcomes in lactating cows following intramammary infection, with 50% of the cows requiring euthanasia after reaching humane endpoints. In contrast, Baker et al. did not observe mortality in lactating cows, likely due to differences in age and lactation stage; Halwe et al. infected older cows late in lactation, while Baker et al. studied younger cows in their first lactation. Our observation of high viral titers in milk post-infection is consistent with findings from both Halwe’s and Baker’s studies. A recent report on intramammary infections of bovine H5N1 in lactating mice found no mortality in either dams or pups^13^. Interestingly, none of the pups tested positive for the virus throughout the study. This suggests that bovine and ferret mammary tissue is more susceptible to bovine H5N1 infection than those of mice.

The mortality rate of H5N1 avian influenza in pregnant women is alarmingly high. A 2008 review documented six cases of pregnant women infected with HPAI H5N1, of whom four succumbed to the infection, while the two survivors experienced spontaneous abortions^14^. Pathological analysis of one fatal case revealed H5N1 viral genomic RNA and antigens in the placenta and fetal lung cells, indicating transplacental transmission of the virus to the fetus^15^. Currently, there are no documented cases of H5N1 infection in lactating women. However, we observed the expression of SAα2,3, the receptor required for H5N1 binding, in human mammary tissue. A limitation of this finding is that our analysis was restricted to non-lactating tissue. Given the expression of the HPAI H5N1 receptor in bovine^16,17^ and ferret mammary glands, as discussed in our manuscript, it is plausible that lactating human mammary tissue may also be susceptible to H5N1 infection.

In conclusion, our study demonstrates the high susceptibility of lactating ferrets and their offspring to HPAI H5N1 infection following intramammary inoculation and milk transmission to suckling newborns. The observed drop in milk production, elevated milk viral titers, and weight loss in ferret dams closely parallel the effects seen in lactating dairy cows after H5N1 inoculation^8,9^. The high neonatal mortality in our study also mirrors the near-total mortality observed in suckling pups during a recent H5N1 outbreak among elephant seals^18,19^, emphasizing the need to understand maternal-neonatal H5N1 infection dynamics across species. These findings underscore the risks of mammary gland infection and milk transmission of H5N1, offering a valuable model for studying viral transmission dynamics and evaluating emerging vaccines and therapeutics against HPAI H5N1.

## Acknowledgements

We thank Dr. Richard Webby (St. Jude Children’s Research Hospital) for providing the initial virus stock and propagation. We thank Dr. Patricia Boley for the technical guidance she provided. We thank Drs. Marilia Cascalho and Jeffery Platt, and Ms. Abigail Williams for critical assessment of our manuscript.

## Materials and Methods

### Animals

Two influenza seronegative pregnant ferrets (*Mustela putorius furo*) were purchased at two weeks prepartum from Triple F Farms (Sayre, PA). Ferret kits were born (n=3/litter) two weeks later and were housed with their nursing mother. Animals were housed in stainless steel cages (Allentown, Inc.) and provided with food and freshwater *ad libitum*. Animals were monitored twice daily, and cages were cleaned once daily.

### Virus

A/bovine/Ohio/B24OSU-342/2024 HPAI H5N1 clade 2.3.4.4b was isolated from samples taken from a dairy farm in the state of Ohio, USA. Samples were sent to St. Jude Children’s Research Hospital for sequencing, passaged twice in eggs and returned to The Ohio State University Plant and Animal Agricultural Research (PAAR) BSL-3 large animal facility. The virus was propagated twice in Madin-Darby canine kidney (MDCK) cells and titrated using plaque assays.

### Intramammary inoculation and ferret monitoring

#### Mammary gland inoculation

Lactating adult female ferrets at 2.5 weeks postpartum were separated from their kits and anesthetized intramuscularly with 0.1 mL dexmedetomidine and maintained with 1-4% isoflurane in 1.5-3 lpm oxygen using a calibrated vaporizer. Animals were weighed, and eye lubricant added. Once anesthetized, the mother was placed on her back, the surrounding area to the nipple was clipped to remove hair and were wiped with cotton pads soaked in 70% ethanol. Milk was expressed manually to determine the location of active mammary glands. A/bovine/Ohio/B24OSU-342/2024 HPAI H5N1 clade 2.3.4.4b (10^5^ EID_50_) in PBS at 125 μL of inoculum was injected via the teat canal into three lactating mammary glands of each ferret. Following inoculation, the ferret dam received 1 mg/kg atipamezole intramuscularly and observed until she was fully awake and returned to the cage and left to nurse suckling neonates.

#### Clinical monitoring

The animals were monitored daily for severity of disease using weight loss and temperature measurements. Mammary glands were also monitored for signs of mastitis (indicated by abnormal milk characteristics, swelling or redness).

### Sample collection and processing

Animals were anesthetized and baseline samples were collected from all animals prior to intramammary inoculation on day 0. Nasal, oral and fecal swabs were collected in a wash buffer (1% BSA and 100 U/ml penicillin, 100 μg/ml streptomycin in PBS) as previously described^12^ and stored at -80°C. Ferret kits had nasal, oral and fecal swabs, weight, temperature, and serum collected on 0, 2, and 4 DPI and ferret dams were sampled on 0, 2, 4, 6 DPI and in addition, milk was collected on those days. Blood was collected from anesthetized animals via the cranial vena cava. Blood was spun twice at 900×g for 10 mins and then at 2,000×g for 15 min. Serum supernatant was collected and stored in -20°C. One kit was humanely euthanized by 4 DPI and BAL fluid, lung, soft palate, and nasal turbinates were collected and preserved in RNAlater (Millipore Sigma) and viral transport media (VTM) (Fisher Scientific) for analysis. Ferret dams were necropsied on 6 DPI and lung, BAL, soft palate, nasal turbinates and mammary gland tissues were collected and stored in both RNAlater and VTM. Tissue samples stored in RNAlater and VTM for viral RNA titration were homogenized using a TissueLyserII (Qiagen), cleared by centrifugation, and supernatant was aliquoted for real-time quantitative PCR (RT-qPCR).

### RNA extraction and RT-qPCR

RNA extraction was performed on nasal, oral, feces, serum, milk, BAL and homogenized tissues. For all samples, 140 μL of sample was used for extraction. All RNA extractions were performed using the QIAamp Viral RNA Extraction kit (Qiagen) following the manufacturer’s recommendations. A one step RT-qPCR (probe based) was carried out using IDT specific Avian Influenza Type A (H5) primer/probe mix (Integrated DNA Technologies) for the identification of influenza A virus clade 2.3.4.4b. Samples were run in duplicate using a protocol of 50°C for 15 min, 95°C for 3 min, and 45 cycles of 95°C for 15 seconds and 60°C for 60 seconds (QuantStudio 3). QuantStudio Design and Analysis software v1.5.3 was used for raw data collection (Thermo Fisher). Data is presented as the average cycle threshold (Ct) value of the duplicates. Undetermined samples were assigned a Ct value of 40.

### Lectin Histochemistry

Sialic acid expression was investigated in lactating ferret mammary gland tissue as well as non-lactating human breast tissue using samples collected from a previously published study (Paquette et al. 2015)^12^. The archived data were generated by sialic acid staining of formalin-fixed, paraffin-embedded mammary tissue sections from healthy lactating ferrets and a commercial human breast cancer microarray panel containing 63 samples, including normal, tumor-adjacent, and tumor tissue. Briefly, tissues were sectioned and mounted to microscope slides and dried on a slide warmer. Tissues were stained with fluorophore-conjugated lectins SNA (*Sambucus nigra*) and MAA (*Maackia amurensis* lectin II) from Vector Laboratories, where SNA is specific for SAα2,6-gal/GalNAc and MAA is specific for SAα2,3-gal-β(1–3)GalNAc. In addition, slides were stained with SYTOX Green nucleic acid stain (Thermo Fisher). Slides were incubated overnight at room temperature. Coverslips were mounted using VECTASHIELD HardSet Antifade mounting medium (Vector Laboratories), and slides imaged on a Zeiss LSM 710 NLO Microscope.

### Statistical analyses

Statistical analyses include two-way analysis of variance (ANOVA) (mixed model) with Tukey’s multiple comparisons test and a log-rank Mantel-Cox test for the survival curve. Statistical significance was defined as *P* ≤ 0.05. All graphs and statistical analyses were performed using GraphPad Prism software version 10.0 (GraphPad Software, San Diego, CA). A limitation of this study is the low sample size (due to animal availability).

